# A high-cholesterol zebrafish diet promotes hypercholesterolemia and fasting-associated liver triglycerides accumulation

**DOI:** 10.1101/2023.11.01.565134

**Authors:** Yang Jin, Darby Kozan, Jennifer L Anderson, Monica Hensley, Meng-Chieh Shen, Jia Wen, Tabea Moll, Hannah Kozan, John F. Rawls, Steven A. Farber

## Abstract

Zebrafish are an ideal model organism to study lipid metabolism and to elucidate the molecular underpinnings of human lipid-associated disorders. In this study, we provide an improved protocol to assay the impact of a high-cholesterol diet (HCD) on zebrafish lipid deposition and lipoprotein regulation. Fish fed HCD developed hypercholesterolemia as indicated by significantly elevated ApoB-containing lipoproteins (ApoB-LP) and increased plasma levels of cholesterol and cholesterol esters. Feeding of the HCD to larvae (8 days followed by a 1 day fast) and adult female fish (2 weeks, followed by 3 days of fasting) was also associated with a fatty liver phenotype that presented as severe hepatic steatosis. The HCD feeding paradigm doubled the levels of liver triacylglycerol (TG), which was striking because our HCD was only supplemented with cholesterol. The accumulated liver TG was unlikely due to increased *de novo* lipogenesis or inhibited β-oxidation since no differentially expressed genes in these pathways were found between the livers of fish fed the HCD versus control diets. However, fasted HCD fish had significantly increased lipogenesis gene *fasn* in adipose tissue and higher free fatty acids (FFA) in plasma. This suggested that elevated dietary cholesterol resulted in lipid accumulation in adipocytes, which supplied more FFA during fasting, promoting hepatic steatosis. In conclusion, our HCD zebrafish protocol represents an effective and reliable approach for studying the temporal characteristics of the physiological and biochemical responses to high levels of dietary cholesterol and provides insights into the mechanisms that may underlie fatty liver disease.

## 1 Background

Cholesterol is an essential sterol lipid and a structural component of cell membranes. Additionally, cholesterol serves as a precursor for steroid hormone and bile salt synthesis and impacts many cell signaling processes (1). Dietary cholesterol is absorbed by intestinal enterocytes, where it is esterified into cholesterol esters (CE) for packaging into chylomicrons, an apolipoprotein B (ApoB) -containing lipoprotein (ApoB-LP), to be transported through the lymphatics and vasculature (2, 3). In circulation, ApoB-LP vary by size and/or tissue of origin (e.g., chylomicrons, very-low-density lipoprotein (VLDL), intermediate-density lipoproteins (IDL), low-density lipoprotein (LDL)). Each ApoB-LP contains one copy of ApoB per lipoprotein particle (4). Cholesterol is also transported by high-density lipoproteins (HDL), which rely on apolipoprotein A1 as a structural component instead of ApoB. While the intestine is the first organ to process dietary cholesterol, the liver also plays a key role in regulating cholesterol homeostasis through a variety of processes: 1) uptake of cholesterol and CE from the circulation from lipoproteins like LDL, 2) conversion of cholesterol into bile acids for excretion, 3) packaging of cholesterol into VLDL for transport to peripheral tissues, and 4) synthesizing cholesterol *de novo* when dietary supplies are limited (5). Dysregulation of cholesterol homeostasis is linked to several lipid disorders including hypercholesterolemia, non-alcoholic fatty liver disease (NAFLD), coronary heart disease, type-2 diabetes, and obesity (6–10).

The zebrafish has emerged as an ideal model for studying human lipid disorders such as hypercholesterolemia and NAFLD (11, 12). The digestive tract of zebrafish and mammals is highly similar and both utilize a similar lipoprotein transport system (13, 14). Furthermore, the small body size, fast growth, and optical transparency make the zebrafish larva optimal for live imaging and high-throughput *in vivo* drug discovery (15). Although traditional rodent models can exhibit elevated hypercholesterolemia in response to high dietary cholesterol, the cholesterol distribution between ApoB-LP and HDL differs between mouse and human (16). One explanation is that mice are naturally deficient in cholesterol ester transport protein (CETP) that transfers cholesterol from HDL to the highly atherogenic ApoB-LP (17, 18). On the other hand, the zebrafish expresses the *cetp* gene and have a more human-like lipoprotein profile than mice (19). A study by Stoletov *et.al.* (20) first showed that feeding HCD to zebrafish caused hypercholesterolemia, lipoprotein oxidation, and vascular lipid deposition. This work suggests that the zebrafish can be a powerful model for studying human lipoprotein regulation and associated lipid disorders.

A recently developed zebrafish ApoB reporter line (LipoGlo) enables the study ApoB-LPs from individual larval zebrafish because it requires 1000-fold smaller amounts of plasma than previous assays (21). LipoGlo contains the Nanoluciferase sequence in the endogenous *apoBb.1* gene locus such that it produces an ApoB-Nanoluc fusion protein enabling the quantification of ApoB-LPs number, size, subcellular distribution and tissue localization (21). Humans express ApoB isoforms that are intestinal-specific (APOB-48) and liver-specific (APOB-100), and both isoforms comprise the profile of ApoB-LPs in a volume of plasma. Zebrafish *apoBb.1* represents 95 % of total *apoB* from both intestine and liver (22, 23). Thus, the LipoGlo zebrafish can be used to measure the quantity, distribution, and sizes of all classes of ApoB-LP (21). Another recently published zebrafish line that facilitates studies of lipid cell biology is the EGFP-*plin2* line which expresses a fluorescent fusion of EGFP to Perilipin 2 (Plin) from the endogenous gene locus (24). As PLINs are evolutionary conserved proteins that coat lipid droplets, this reporter line allows direct monitoring and quantification of the number and size of lipid droplets. Together the ApoB and Plin reporter lines enable us to track both intracellular and plasma lipids providing opportunities to evaluate the effect of HCD on zebrafish lipid physiology.

Different standard diets containing various combinations of cholesterol and other nutrients, such as carbohydrates and protein, have been extensively tested in murine models (25). Although custom-made HCD has been previously used for studying lipid metabolism in zebrafish (20, 26), there has yet to be a uniformly adopted protocol throughout the zebrafish field, and the effect of HCD in the zebrafish has not been thoroughly evaluated. In this study, we employed an improved HCD protocol for zebrafish, replacing the ether solvent used to solubilize the cholesterol with less toxic ethanol in the preparation of the HCD. Our primary goal of this study was to address how dietary cholesterol alters lipoprotein regulation and lipid deposition in zebrafish. We found that the HCD increased ApoB-LP levels in a dose- and time-dependent manner. More interestingly, we have identified a fasting-dependent increase in liver triglyceride (hepatic steatosis) in zebrafish fed an HCD, suggesting a novel mechanism for cholesterol-induced triglyceride accumulation in the liver. We have also uncovered increased expression of the *fatty acid synthase* (*fasn*) gene expression in adipose tissue of the HCD zebrafish, which suggests that fasting-dependent hepatic steatosis is associated with *de-novo* lipogenesis in adipose.

## 2 Methods

### 2.1 Ethics statement

All procedures using zebrafish were approved by the Carnegie Institution Department of Embryology Animal Care and Use Committee (Protocol #139).

### 2.2 Zebrafish husbandry and maintenance

All zebrafish (*Danio rerio*) were raised and maintained in the zebrafish facility at the Carnegie Institution for Science, Baltimore, MD, United States. Zebrafish adults were maintained at 27a°C on a 14:10 h light:dark cycle and fed once daily with ∼3.5 % body weight Gemma Micro 500 (Skretting USA). Embryos were collected by natural spawning and raised in a 28.5°C incubator with 14:10 h light:dark cycle before moving to the fish facility for initial feeding at 6-day post fertilization (dpf). Unless otherwise specified, the fish larvae were initially fed with GEMMA Micro 75 two times daily until 14 dpf. Juvenile (15-42 dpf) were fed with GEMMA Micro 150 twice and *Artemia* once daily. Fish older than 42dpf were fed with GEMMA Micro 300 once daily.

### 2.3 Diet preparation

The 1 %, 2 %, 4 %, and 8 % cholesterol diet was made by dissolving 10 mg, 20 mg, 40 mg and 80 mg of cholesterol (C8667, Sigma-Aldrich, St. Louis, MO, United States) respectively in 10mL 100 % ethanol (Figure 1 A). After the cholesterol was fully dissolved, 990 mg, 980 mg, 960 mg and 920 mg of zebrafish diet (GEMMA, Skretting AS, Westbrook, Maine, United States), respectively, was added. The mixture was shaken on a horizontal shaker in a fume hood until all visible liquid was evaporated, followed by leaving in the fume hood overnight ensuring that ethanol was fully evaporated. Dried HCD diet was pushed through a sieve (50-100 µm for G75, 150 µm for G150, and 500 µm for G500) using a pestle and stored at -20D. The control diet was treated the same as the HCD, except no cholesterol was added. Fresh diets were made for each feeding trial.

**Figure 1.**
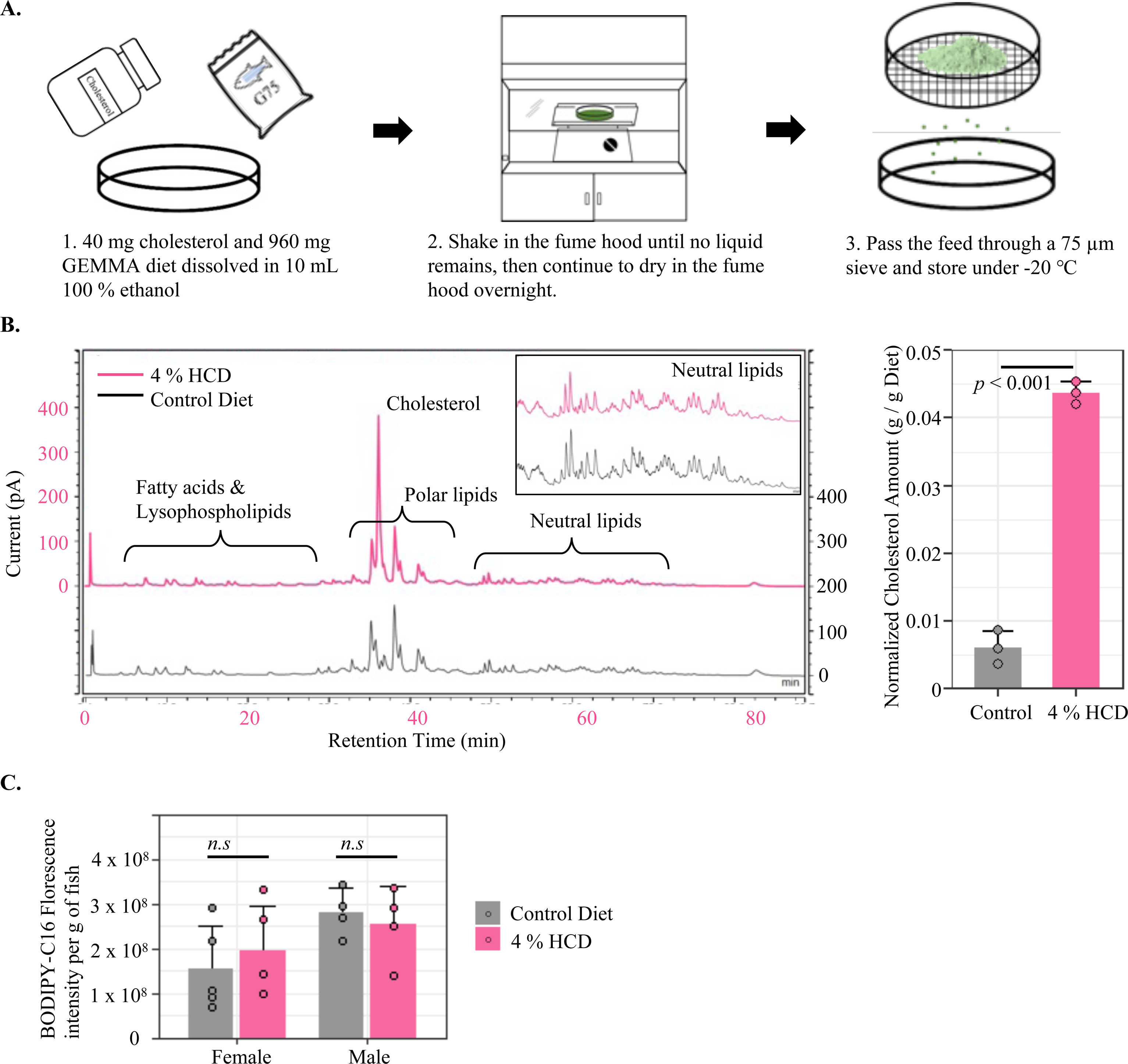
Diet-making protocol and quality control. **A)** Procedure for making the HCD. **B)** Quantification of lipids in diets using HPLC-CAD. The left figure shows representative chromatographs for extracted lipids from an HCD and the control diet. The bar graph on the right compares the cholesterol amount between HCD and the control diet (n=3, mean +/- SD, Welch t-test). **C)** Comparison of food intake between fish fed HCD and control diet with 0.025 % BODIPY-C16. Total fluorescence was measured from lipids extracted from the entire intestinal tract and liver of fish after 1 h of feeding (n > 4, mean +/- SD, Two-Way ANOVA with TukeyHSD post-hoc tests, n.s *p* > 0.05).

### 2.4 Feeding trials

All larval feeding trials were performed in a 3 L tank with 60 zebrafish larvae. Animals were fed either a control diet or an HCD from initial feeding at 6 dpf to 14 dpf. A time- and dose-response experiment was performed by feeding LipoGlo (*apoBb.1^NLuc/NLuc^*) (21) zebrafish either the control diet, 1 %, 2 %, 4 %, or 8 % HCD from initial feeding at 6 dpf to 14 dpf. Fish were sampled 2 h after the morning feeding at 6 dpf (after initial meal), 7 dpf, 9 dpf and 14 dpf. Two replicate tanks were used for each group. All sampled fish were anesthetized with tricaine, then placed under a stereo microscope to ensure the fish had food in their intestines. Four fish from each tank were snap-frozen on dry ice, and then kept at -80°C until further LipoGlo-counting (21) was performed to measure total ApoB-LP concentration. Five fish from each tank were immediately imaged under a Nikon SMZ1500 microscope with an HR Plan Apo 1x WD 54 objective, Infinity 3 Lumenera camera and Infinity Analyze 6.5 software. All fish were imaged under transmitted light with the same exposure and white balance settings. The experiment was repeated 3 times with 3 different clutches of animals obtained from natural crosses of LipoGlo adults.

A fasting experiment was done by feeding fish either control diet or 4% HCD from 6 dpf to 13 dpf. After their last meal, fish from two replicate tanks were merged and randomly split into two new tanks. One tank was fed the same HCD or control diet for another day, while the other tank fasted for 24 h or 48 h. The experiment was repeated 3 times using 3 different clutches of fish. One clutch of fish was fasted for 72 h. All fish from each day were imaged as described above.

Another fasting experiment was performed by initially feeding fish either control diet or 4 % HCD from 6 dpf to 13 dpf. Six fish from each dietary group were transferred and fasted in a 12-well cell culture plate with 1 fish per well. The same individual fish were repeatedly imaged 0 h, 15 h, 24 h and 48 h after last meal using an Axiozoom V16 microscope equipped with a Zeiss Plan NeoFluar Z 1x/0.25 FWD 56 mm objective, AxioCam MRm camera, and Zen 2.5 software. The experiment was repeated 2 times with 2 different clutches of fish.

The adult feeding trial was conducted in 3 L tanks with no more than 30 fish per tank. One-year old fish were distributed equally into two 3 L tanks, with equal gender per tank. Fish were fed either the control diet or HCD for 2 weeks. Postprandial fish were sampled 2 h after the last meal while fasting fish further fasted for 3 days. Sampled fish were anesthetized with tricaine and dissected with a clean cut between the anus and caudal peduncle. Blood samples were taken using EDTA coated Kunststoff Kapillaren tubes (Sanguis Counting, Nümbrecht, Germany) and then transferred to 1.5 mL Eppendorf tubes. After centrifuging at 4 °C for 2 - 5 min at 5000 g, 1 - 3 µL of plasma were transferred into a new 1.5 mL Eppendorf tube and snap-frozen on dry ice. The middle intestine (27, 28) and liver were dissected and snap-frozen on dry ice. A scalpel was used to take off the white skeletal tail muscle between the region of pelvic fins to the caudal part. Fish skin was carefully removed, and only upper half of the muscle was kept for further analysis. All samples were stored under -80°C before further analysis.

To compare the food intake between the 1-year-old fish fed the HCD and control diet, we used a fluorescent fatty acid (BODIPY-C16) as a dietary tracer and measured total fluorescent levels in the entire digestive tract (29). This was done by adding 0.5 mg (0.05 %) of BODIPY-C16 in 1 g of the control diet or 4 % HCD diet. Two tanks of fish, each with 5 females and 4 males, were fed either the control diet or 4 % HCD containing 0.05 % BODIPY-C16. The same protocol was used to prepare the diets except that BODIPY-C16 was added together with cholesterol during the process. A total amount of 0.315 g BODIPY-C16 diet was given to each tank, which was estimated as 3.5 % of fish weight. After 2 h of feeding, the entire intestine with diet in the lumen was carefully dissected. Total lipids were extracted immediately for fluorescent measurement as previously described (29).

### 2.5 LipoGlo Assays

The generation of *apoBb.1^NLuc/Nluc^* fish and analytical protocols (LipoGlo Assays) were previously described (21). In brief, TALEN technology was used to fuse zebrafish *apoBb.1* gene with a cDNA sequence encoding Nanoluc^®^ luciferase. These fish (*apoBb.1^Nluc/Nluc^* or *^Nluc/+^* for the fusion protein) were used as described (21).

Sampled whole LipoGlo larvae or pieces of LipoGlo adult tissue were placed in individual wells of a 96-well plate and filled with 100 µL ApoB-LP stabilization buffer (21). Fish were then homogenized in a microplate-horn sonicator (Qsonica, Q700 sonicator with 431MPX microplate-horn assembly) filled with 17 mm of chilled, reverse osmosis (RO) water and processed at 100 % power for a total of 30 s, delivered as 2 s pulses interspersed with 1 s pauses. Homogenates were stored on ice for immediate use, or frozen at −80 °C and then thawed on ice for later use. Quantification of ApoB-LP levels (LipoGlo-Counting) was prepared by mixing 40 µL homogenate with an equal volume Nanoluc reaction buffer (PBS: NanoGlo Buffer: NanoGlo substrate furimazine, 30: 10: 0.2) in a black perkin elmer plate (6065400, PerkinElmer, Waltham, MA, USA). Within 2 min of buffer addition, the plate was read under a SpectraMax M5 plate reader (Molecular Devices, San Jose, CA, USA) with top-read chemiluminescent detection and a 500 ms integration time.

Three homogenates from each group were selected for LipoGlo-Electrophoresis, which was used to quantify the ApoB-LP size distribution as previously described (21). A 3 % native polyacrylamide gel was prepared and cast overnight at 4 °C using a 1 mm spacer plate and 10-well comb. The next day, the gel was put into a mini-protean electrophoresis rig filled with pre-chilled 1x TBE at 4 °C and pre-run at 50 V for 30 min. Afterwards, 12.5 µL of sample solution (sample homegenate: 5x loading dye, 4: 1) was loaded per well. DiI-labeled human LDL (L3482, Thermo Fisher Scientific) solution (DiI-LDL: 5x loading dye, 4: 1) was used as a migration standard. The gel was run at 50 V for 30 min, followed by 125 V for 2 h. After the run, the gel was first equilibrated in an imaging solution (1 mL + 2 µL NanoGlo substrate) for 3 min, and then placed into an Odyssey Fc (LICOR Biosciences) gel imaging system and imaged in the chemiluminescence channel for 2 min (NanoLuc detection) and then the 600 nm channel for 30 s (DiI LDL standard detection). The imaged gel was quantified using ImageJ based on the previously described method (21).

To visualize the whole-organism localization of ApoB-LP (LipoGlo-Microscopy), 14 dpf zebrafish larvae were anesthetized and fixed in 4 % paraformaldehyde (PFA) for 3 h at room temperature and rinsed 3 times for 15 min each in PBS-Tween (PBS containing 0.1 % Tween-20 detergent). Fixed larvae were then mounted in 50 µL low melting point agarose (0.01 g/mL TBE) containing Nano-Glo substrate (1 %). Nanoluc images were taken by Zeiss Plan NeoFluar Z 1x/0.25 FWD 56 mm objective, AxioCam MRm camera, and Zen 2.5 software (21).

### 2.6 Liver EGFP-Plin2 confocal imaging

We used the previously developed EGFP-Plin2 reporter line (24) to directly monitor and quantify the number and size of lipid droplets in zebrafish larvae. *Fus(EGFP-plin2)*/+ larvae were fed either 4 % HCD or control diet from 6 dpf to 13 dpf, followed by fasting for 24 h. Confocal images of the liver were obtained from 4 fish fed HCD and 4 fish fed the control diet. Images were taken on a Leica SP5II confocal microscope with a 63×1.4 HCX PL Apo oil immersion lens.

### 2.7 Feeding trial on fabp6-GFP reporter line

To examine the effect of dietary cholesterol on zebrafish bile metabolism, we fed the fish HCD and control diets and bile production was examined. A previously established *Tg(-1.7fabp6: GFP)* reporter line expresses green fluorescent protein (GFP) in the ileal epithelium driven by a 1.7-kb zebrafish *fabp6* promoter fragment was used in the feeding trial (30). This reporter line can be used to monitor bile salt signaling in zebrafish because bile salts bind and activate farnesoid X receptor (FXR) which in required to drive expression of *fabp6: GFP* (30). The feeding trial was performed in the fish facility in Duke University, Durham, NC, United States following protocols approved by the Duke University Medical Center Institutional Animal Care and Use Committee (protocol numbers A115-16-05 and A096-19-04). Since Cyp7a1 is the first and rate limiting step in bile salt synthesis pathway, we used transgenic mutant fish *cyp7a1^-/-^; Tg(-1.7fabp6: GFP)/+* and wild-type *cyp7a1^+/+^; Tg(-1.7fabp6: GFP)/+* (30) to monitor bile salt production. The fish were fed either HCD or control diet from initial feeding at 6 dpf to 13 dpf, followed by fasting 24 h. The fluorescence in the transgenic reporter line was quantified using a Leica M205 FA stereomicroscope with exposure time and magnification as described previously (30).

### 2.8 HPLC

The quantification of specific fish lipids or lipid classes was performed by using high performance liquid chromatography system with a Dionex Corona Veo charged aerosol detector (HPLC-CAD from Thermo Fisher Scientific, Waltham, MA, USA). The HPLC-CAD and analytical methods were previously described (31). Lipids were extracted from individual tissue homogenates using the Bligh-Dyer method (32), then dried and resuspended in HPLC-grade isopropanol. Lipid components of each sample were detected by the HPLC system with Accucore C18 column (150 × 3.0 mm, 2.6 µm particle size) and a CAD. A gradient mobile phase was used for separating different lipid classes: 0 - 5 min = 0.8 mL/min in 98 % mobile phase A (methanol-water-acetic acid, 750:250:4) and 2 % mobile phase B (acetonitrile-acetic acid, 1000:4); 5 - 35 min = 0.8 - 1.0 mL/min, 98 - 30 % A, 2 - 65 % B, and 0 - 5 % mobile phase C (2-propanol); 35 - 45 min = 1.0 mL/min, 30 - 0 % A, 65 - 95 % B, and 5 % C; 45 - 73 min = 1.0 mL/min, 95 - 60 % B and 5 - 40 % C; and 73 - 80 min = 1.0 mL/min, 60 % B, and 40 % C.

To further identify peaks from different lipid classes on HPLC, thin-layer-chromatography (TLC) was first used to separate lipid classes from total lipid extracts of tissue homogenates (33). Total lipid extracts were dried and resuspended in 50 µL isopropanol, after which 20 µL were loaded in one row of a 20 cm × 20 cm channeled silica plate (LK5D, Whatman, Maidstone, United Kingdom) and another 20 µL were loaded on the adjacent channel. Lipid standards of triolein, cholesterol oleate, phosphatidylcholine (PC), and phosphatidylethanolamine (PE) (all from Sigma-Aldrich) were loaded on the adjacent columns. The plate was first run in the polar solvent (ethanol: triethylamine: ddH2O: CHCl3, 27: 25: 6.4: 25) to halfway up of the plate (10 cm), air-dried and then run in non-polar solvent (petroleum ether: ethyl ether: acetic acid 64: 8: 0.8) to near the top of the plate. The whole plate was then air-dried and cut in half. One half of the plate with one replicate of each sample and standards was sprayed with chromic-sulfuric acid (5 % w/v potassium dichromate solution in 40 % v/v aqueous sulfuric acid solution) and placed in oven (> 180 °C) to char the lipids. After cooling down, the lipid could be visualized and identified based on the position of the lipid standards. Based on the position of lipid bands on the charred plate, the lipid bands of TG, CE, PC, and PE from the uncharred plate were separately scraped out into 10 mL glass tubes for lipid extraction. One milliliter of chloroform: methanol (2:1) was added into the tube with silica, which was vortexed and centrifuged for 5 min at 2000 g. Afterward, 0.8 mL upper liquid was carefully transferred into a new tube, and the whole process was repeated 3 times. The collected 2.4 mL chloroform: methanol (2:1) was centrifuged for 5 min at 2000 g again to ensure all silica was sedimented at the bottom. Two milliliters of upper liquid were carefully collected, dried under N2, resuspended in 50 µL isopropanol, and then subject to HPLC-CAD as described above.

### 2.9 RNA isolation, quantitative RT-PCR and transcriptomic sequencing (RNA-seq)

Total RNA was extracted from the livers or adipose tissue using Direct-zol RNA Microprep Kits (R2061, Zymo Research, Irvine, CA, USA), according to the manufacturer’s instructions. The RNA concentration and integrity were measured by a Nanodrop One (Thermo Fisher Scientific) and a Bioanalyzer (Agilent Technologies, Santa Clara, USA). All extracted RNA had integrity (RIN) values higher than 8, which indicates sufficient RNA quality for sequencing. The cDNA sequencing libraries were prepared using a TruSeq Stranded mRNA Library Prep Kit (Illumina, San Diego, USA) according to the manufacturer’s instructions. Libraries were sequenced using 75 bp single-end mRNA sequencing (RNA-seq) on Illumina MiSeq (Illumina, San Diego, CA, USA) at the Department of Embryology, Carnegie Institution for Science (Baltimore, MD, USA). An average of 40 million reads was acquired from each sample. The reads were aligned to the zebrafish genome (GRCz11) using STAR (34). The resulting .bam files were subsequently used to generate raw gene counts using *featureCounts* (35). Ensembl Gene IDs were used to identify genes in this study. The raw fastq files are publicly available on Sequence Read Archive (SRA) with accession number PRJNA998935.

For quantitative RT-PCR, the cDNA was synthesized using the iScript cDNA Synthesis Kit (Bio-Rad Laboratories, Inc, 1708891). All cDNA samples were then prepared using SsoAdvanced Universal SYBR Green Supermix (Bio-Rad Laboratories, Inc, 1725271). The primer pairs targeting zebrafish *fasn* transcripts (forward: 5’-CTGGCCATGGTCCTTAAAGATGG-3’, reverse: 5’-AGTTGGCGAAGCCGTAGTTG-3’) were used for gene expression analysis and zebrafish 18 S gene (forward: 5’ - GAACGCCACTTGTCCCTCTA - 3’, reverse: 5’-GTTGGTGGAGCGATTTGTCT - 3’) was used as the reference gene. The quantitative RT-PCR was performed in triplicate for each sample with the Bio-Rad CFX96 Real-Time System with 45 cycles: 95 °C for 15 s, 59 °C for 20 s, and 72 °C for 20 s. All results were analyzed with the Bio-Rad CFX Manager 3.0 software and relative gene expression was calculated using the ΔΔCT method (36).

### 2.10 Statistics

Differential expression analysis (DEA) was performed using R (v.4.1.2) package edgeR (37). Only genes with a minimum count level of at least 1 count per million (CPM) in more than 50 % of samples from each tissue were kept for further DEA. A generalized linear model (GLM) approach described in the edgeR manual was used to find differentially expressed genes (DEGs) between HCD and control groups in each sex separately. DEGs were determined if a gene had a false discovery rate (FDR) adjusted *p* value (*q* value) < 0.05 and absolute log2 fold change (|Log2FC|) > 1.

All statistical analyses were performed using R v.4.1.2. All datasets were first subjected to Shapiro-Wilk’s test for normality. One-way or two-way analysis of variance (ANOVA) was used for testing the main effect, and Tukey’s honestly significant difference (HSD) was then used for post hoc testing. If the data was not normally distributed, Welch’s ANOVA from R package WRS2 was used for testing the main effect and the Games-Howell method for post-hoc testing.

## 3 Results

### 3.1 Generation of a high-cholesterol diet

We developed an improved high-cholesterol diet (HCD) paradigm for zebrafish, which can be used for studying the impact of dietary lipids on lipoprotein metabolism and overall digestive organ physiology (Figure 1A). We used a popular commercially available diet (Gemma, Skretting, Norway) as our basal diet, which contains 14 % lipids and 59 % protein (https://zebrafish.skrettingusa.com/). Since Gemma is not an open-source formulation, we analyzed its lipid profile and found the predominate lipids were phospholipids, and the basal cholesterol level was about 0.6 % of the diet (Figure 1B). A 4 % HCD was made by dissolving 0.04 g of cholesterol in 100 % ethanol before adding 0.96 g of Gemma. This mixture was shaken in a fume hood until no liquid remained and further dried overnight in the fume hood. The dried food was ground through sieves (50-100 µm for G75, 150 µm for G150, and 500 µm for G500) to make the final HCD product (Figure 1A). By analyzing the lipid profiles of the diet, we found that the 4 % HCD contained an average of 4.3 % of cholesterol in the diet, compared to 0.6 % cholesterol in the control diet (Figure 1 B). This indicated that the HCD-making process is highly effective and reproducible as there was little variation between different batches. All other phospholipid and neutral peaks were identical between the control diet and 4 % HCD (Figure 1 B).

To further evaluate the effect of dietary cholesterol and ethanol treatment on fish palatability, we used fluorescent fatty acid BODIPY-C16 (0.025 % of feed) as a dietary tracer and measured total fluorescent levels in the entire digestive tract after a single meal (29). Adult zebrafish were fed either the control diet or 4 % HCD, and whole guts, including gut contents, were dissected for total lipid extract florescence. Fish fed either diet had similar levels of gut lipid fluorescence, indicating that the 4 % HCD was equivalently palatable as the standard feed (Figure 1 C).

### 3.2 Feeding HCD increases total ApoB-LP abundance in zebrafish larvae

To examine the effects of dietary cholesterol on ApoB-LP zebrafish physiology, we fed HCD to the LipoGlo fish through 14 dpf to monitor and quantify ApoB-LP using LipoGlo assays as previously described (21) (Figure 2 A). We found juvenile animals (14 dpf) fed a 4 % HCD had increased overall ApoB-LP levels throughout the body (Figure 2 B, 24 fish from each diet group). To determine if the HCD diets altered developmental growth rate, total standard length was measured (38). Fish fed 4 % HCD had similar standard length (5.13 ± 0.51 mm) as compared to fish fed the control diet (5.20 ± 0.47 mm).

**Figure 2.**
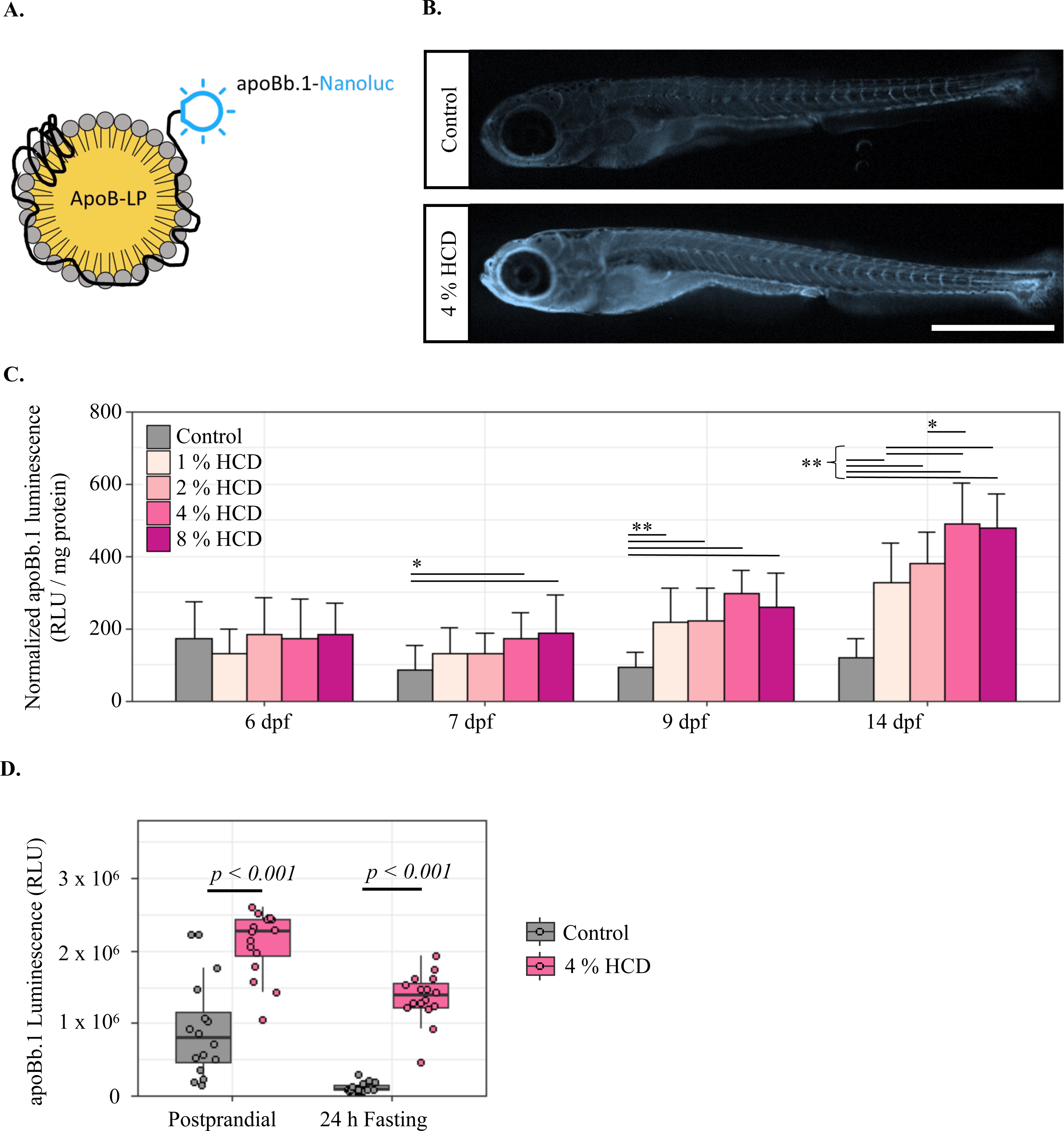
Effect of HCD on ApoB levels in zebrafish larvae. **A)** The LipoGlo reporter line, which contains an engineered *apoBb.1* gene fused with a nanoluciferase reporter was used for quantifying ApoB-LP levels in fish (21). **B)** LipoGlo-imaging was used to monitor ApoB-nluc chemiluminescence in whole zebrafish larvae (14 dpf) fed either the control diet or 4 % HCD. 23 fish were sampled from each dietary group and a representative image is shown. Scale = 1 mm. **C)** LipoGlo-counting quantified whole-body ApoB-LP levels in each diet from initial feeding at 6 - 14 dpf. Each bar represents mean ± SD from 24 fish (robust Two-Way ANOVA with Games-Howell tests). * *p* < 0.05, ** *p* < 0.001 **D)** Comparison of ApoB-LP levels (LipoGlo-counting) in fish fed the HCD and control diet at postprandial or 24h-fasted stage (n =16 fish each group, robust Two-Way ANOVA with Games-Howell tests).

To further confirm that the increased ApoB-LP signals was indeed caused by the HCD, we fed *apoBb.1^Nluc/+^* transgenic animals diets containing various concentrations of cholesterol (0 %, 1 %, 2 %, 4 %, and 8 %) and quantified ApoB-LP (RLU per mg of protein) in whole zebrafish lysates from 6 to 14 dpf. After one day of feeding, at 7 dpf, the fish under the conditions of 4 % and 8 % HCD exhibited significantly (*p* < 0.05) higher ApoB-LP levels compared to fish fed the control diet (Figure 2 C). The ApoB-LP levels in fish fed 1 % and 2 % HCD appeared to increase later and were significantly (*p* < 0.05) higher than the control diet from 9 dpf onwards. Furthermore, 14 dpf fish fed 4 % and 8 % HCD had significantly (*p* < 0.05) higher levels of ApoB-LP compared to fish fed 1 % and 2 % HCD (Figure 2 C). A 24 h-fast at 13 dpf significantly (*p* = 0.001, robust 2-way ANOVA) decreased ApoB-LP levels regardless of dietary cholesterol levels, and the levels of ApoB-LP remained significantly higher (*p* = 2 × 10^-5^, Games-Howell test) in fasted fish fed HCD compared to control diet (Figure 2 D).

A HCD protocol which used diethyl ether to deliver extra cholesterol to a standard fish diet was previously published (20). As diethyl ether is toxic to animals by inhalation or ingestion, we have improved the method by using more readily available and less hazardous ethanol to make the HCD. The efficiency of the two protocols was evaluated by comparing the dietary effect of HCD on ApoB-LP levels and indicated that ether- or ethanol-made HCD was able to increase the ApoB-LP levels in zebrafish larvae (14 dpf; Supplementary Figure 3). We also tested if vacuum treatment could improve the cholesterol-delivering efficiency of the HCD; however, no difference in ApoB-LP levels between the fish fed untreated and vacuum-treated HCD was observed (Supplementary Figure 3).

### 3.3 Fasting after an HCD leads to liver steatosis

Larval zebrafish fed the 4 % HCD from 6 dpf to 13 dpf then fasted for 24 h developed an opaque liver phenotype (dark under transmitted light; Figure 3A, n > 20 fish from each diet group). After staining the liver with Oil Red O, fish exposed to the HCD showed more staining, indicating higher neutral lipid accumulation in vasculature and liver (Figure 3 B, 40 fish from each diet group). Since PLINs are evolutionally conserved proteins that coat lipid droplets, the *EGFP-plin2* reporter line allows us to visualize lipid droplets in live zebrafish at the tissue and subcellular level (24, 39). Therefore, we fed 4 % HCD to *Fus(EGFP-plin2)/+* fish to visualize the subcellular location of the accumulated lipids (24). Indeed, more lipid droplets were observed in the liver of fish fed 4 % HCD compared to the control diet (Figure 3 C).

**Figure 3.**
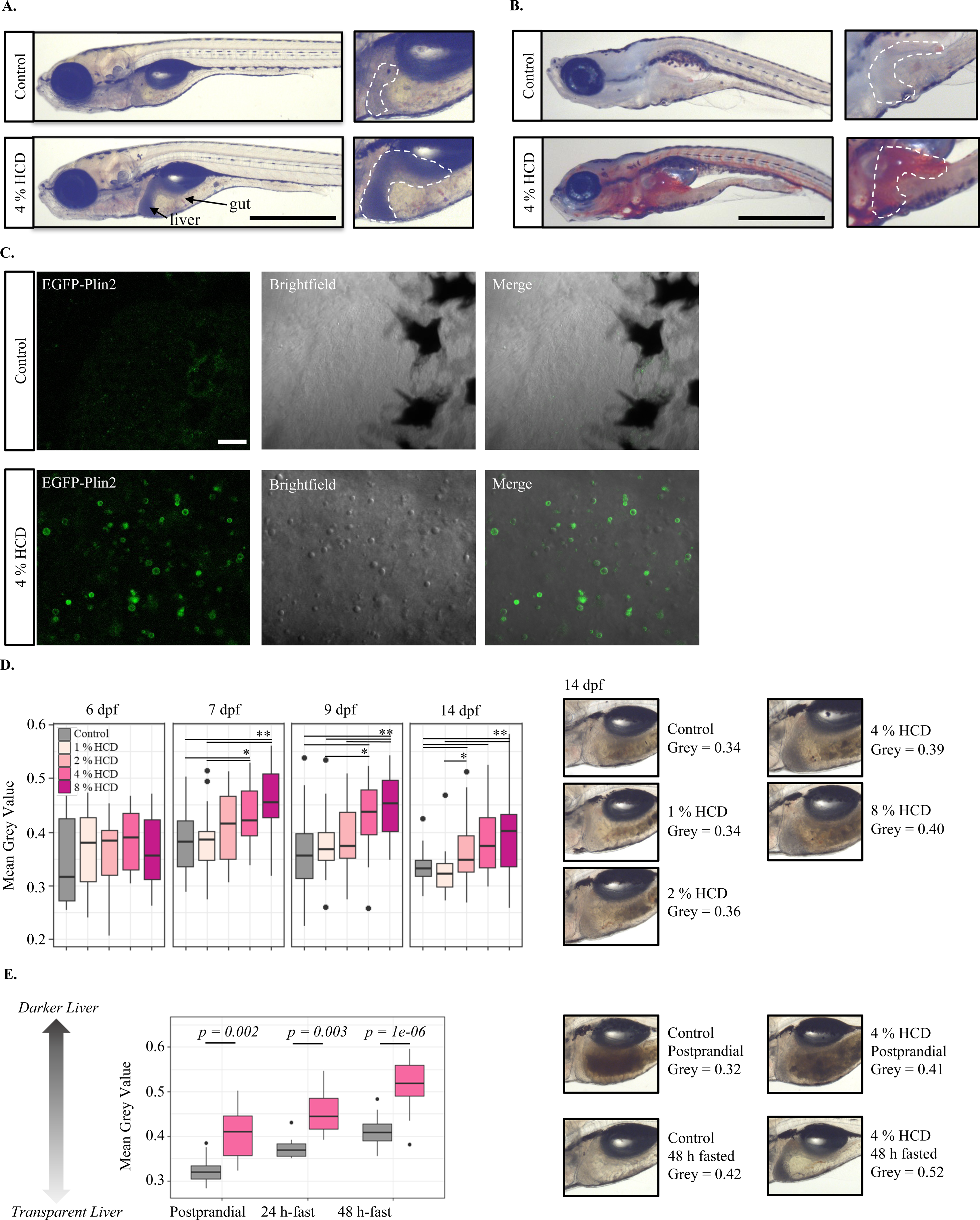
Effect of HCD on liver opacity of zebrafish larvae. **A)** Representative brightfield image of 14 dpf zebrafish larvae fed the control diet or 4 % HCD from 6 dpf to 13 dpf followed by 24h fasting (n > 20 fish per group). Scale = 1 mm. **B)** Representative Oil-red-O staining image of 14 dpf zebrafish larvae fed the control diet or 4 % HCD from 6 dpf to 13 dpf followed by 24h fasting (n = 38 fish per group). Scale = 1 mm. **C)** Confocal images of the liver of EGFP-Plin2 zebrafish, which express EGFP coding sequence fused to the N-terminus of the endogenous *plin2* gene. Representative images were taken from the livers of 14 dpf fish fed 4 % HCD or control diet (n = 4 fish per group). Scale = 10 µm. **D)** Measurement of liver opacity in postprandial fish (2 h after latest meal) fed control diet, 1 %, 2 %, 4 %, or 8 % HCD. Two hundred individuals were used in total. Liver opacity was calculated by measuring the mean grey value of the marked liver area (representative figures shown on the right), which were then normalized by the mean grey value of the muscle area next to the liver. A robust Two-Way ANOVA with Games-Howell tests was used for statistical test. Details of the method are illustrated in Supplementary Figure 4. **E)** Effect of fasting on liver opacity. Fish were fed either the control diet or 4 % HCD from 6 dpf to 13 dpf, followed by either continued feeding or fasting for 24 h or 48 h. Liver opacity was calculated using the same method as Figure 3 D. A robust Two-Way ANOVA with Games-Howell tests was used for statistical test. Figures of the other two experiments are shown in Supplementary Figure 5. * *p* < 0.05, ** *p* < 0.001

To further characterize the liver steatosis observed in animals fed the HCD, we measured the liver opacity in postprandial fish fed various cholesterol concentrations (0 %, 1 %, 2 %, 4 %, and 8 %). The opacity of the liver was quantified using ImageJ to measure the mean grey value of the entire liver area, which was then normalized by the grey value of an adjacent area (Supplementary Figure 3). Feeding of 4 % and 8 % HCD significantly (*p* < 0.05) increased the liver opacity from 7 dpf fish (Figure 3 D, n = 10 for each dietary group). No liver opacity differences were observed in fish fed the control diet, 1 %, and 2 % HCD, except that 14 dpf fish sometimes had a darker liver when fed 2 % HCD than 1 % HCD (*p* = 0.03, Figure 3 D). However, considerable variation was observed in the opacities of the livers between both individuals and different strains, for example, 30 % (9 out of 30 fish) of the liver remained transparent in 14 dpf fish fed feeding 8 % HCD. All fish were fed twice daily (8:00 in the morning and 13:00 in the afternoon). Since there is a 19h interval between each afternoon meal and the next day morning meal, we hypothesize that fasting is another factor that influences liver opacity.

Therefore, an experiment was conducted for monitoring and quantifying liver steatosis associated with the degree of fasting. Fish were fed 4 % HCD or control diet from 6 dpf to 13 dpf followed by fasting for 24 h or 48 h, while postprandial fish were fed the same diet up to 14 dpf. Although the liver opacity is significantly (*p* < 0.05) different between fed HCD and the control diet regardless of fasting, the level of differences was much larger in the 48 h-fasting group (*p* = 1 × 10^-6^, coefficient of variation = 0.1) than the postprandial group (*p* = 0.002, coefficient of variation = 0.14) (Figure 3 E). By visualizing liver opacity under the microscope, we observed 100 % dark liver (n = 19) in HCD-fed fish after 48 h fasting, while only 75 % of the 24 h-fasted fish (9 out of 12 fish) and 64 % of the postprandial HCD-fed fish (9 out of 14 fish) had dark livers. The fasting experiment was repeated 3 times with 3 fish stocks, and we observed the same results (Supplementary Figure 5).

Another fasting experiment was conducted to track liver steatosis in individual fish during fasting. Individual fish were imaged repeatedly at 0 h, 15 h, 24 h, and 48 h after fasting (n = 6 for each dietary group). A similar result was that the difference in liver opacity became larger after 48 h of fasting (*p* = 0.0007) than in postprandial fish (*p* = 0.009, Supplementary Figure 6). The experiment was repeated twice with two separate fish stocks. After 48 h of fasting, 92 % (11 out of 12) of fish on HCD developed dark liver, while the phenotype was not observed in control fish (Supplementary Figure 6). Both fasting experiments have suggested that fasting is highly associated with liver opacity.

### 3.4 Feeding HCD followed by fasting causes hypercholesterolemia and hepatic steatosis in adult zebrafish

To further investigate the effect of the HCD on the tissue and organ level, we have conducted an HCD feeding trial on one-year-old zebrafish adults. Zebrafish develop white adipose tissue which can store increasingly large amounts of neutral lipid as animals increase in size (40). Neutral lipid stored in adipose tissue in juvenile or adult zebrafish can be mobilized by fasting and eventually depleted as early as 7 days of fasting (41–43). Therefore, we speculated that adult fish under HCD needed more than 3 days of fasting to develop visible hepatic steatosis. We fed adult zebrafish 4 % HCD or the control diet for two weeks, followed by a 3-day fast.

No significant difference was observed for plasma ApoB-LP levels between control- and HCD-fed postprandial fish, though HCD female had slightly higher values on average (2 × 10^6^ *vs.* 1.6 × 10^6^ RLU/µL plasma, Figure 4 A). After fasting for 3 days, the HCD fish had significantly increased the ApoB-LP levels in plasma (*p* < 0.001 in females and *p* = 0.02 in males), while ApoB-LP levels in the fasted control fish remained the same as the postprandial controls (Figure 4 B). We subjected 3 female plasma samples from each dietary group to LipoGlo electrophoresis to characterize the ApoB-LP sizes (zero mobility (ZM), VLDL, IDL, and LDL) based on their migration distance (21). Although postprandial fish fed the 4 % HCD had significantly (*p* < 0.05) higher VLDL, IDL, and LDL, the percentage of each ApoB-LP class remained the same between the HCD and control fish (Figure 4 A). The increase of ApoB-LP levels in female plasma reflects increased VLDL and ZM classes, while the IDL and LDL levels were similar between fish fed HCD and control diet (Figure 4 B).

**Figure 4.**
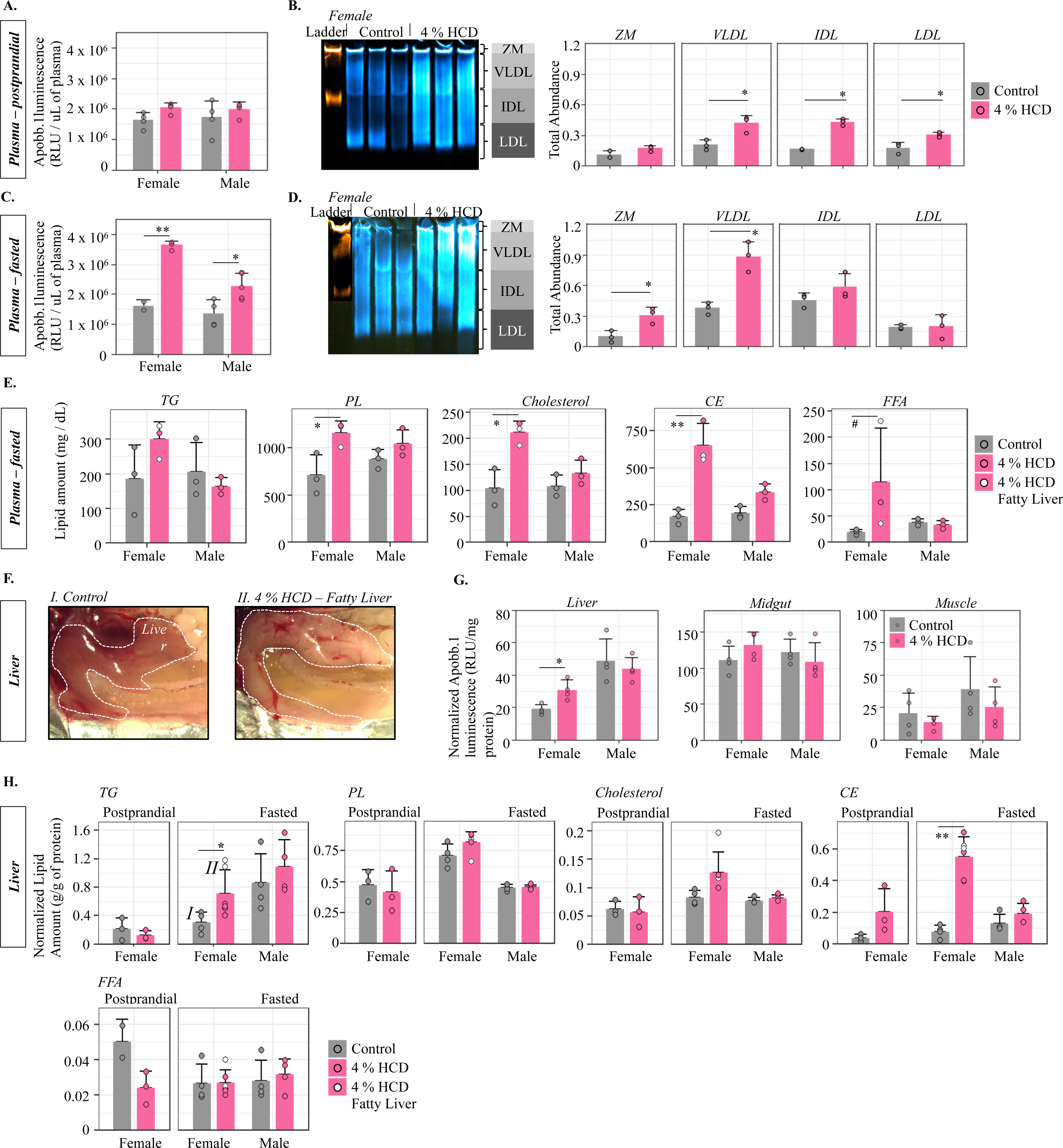
Effect of HCD on plasma ApoB-LP levels and plasma and liver lipids compositions of adult fish. **A)** LipoGlo-counting for measuring total ApoB-LP in plasma of postprandial fish fed the control diet or 4 % HCD for 2 weeks (mean +/- SD, Two-Way ANOVA with Tukey’s post-hoc tests). **B)** The Gel figure is LipoGlo-Electrophoresis (a Native-PAGE assay to separate the ApoB-LP based on size) on 3 representative female plasmas from each diet (mean +/- SD, student t-tests). **C)** LipoGlo-Counting (mean +/- SD, Two-Way ANOVA with Tukey’s post-hoc tests) and (D) LipoGlo-Electrophoresis (mean +/- SD, student t-tests) on plasma samples from fish fed the control diet and 4 % HCD, followed by a 3-day fasting period. **E)** Quantification of major lipid classes in fasted zebrafish plasma using HPLC-CAD platform. Zebrafish that were identified to carry the “fatty liver” phenotype are indicated as white dots on the bar graphs. A robust Two-Way ANOVA with Games-Howell tests was used for statistical test. **F)** Representative liver images from fish fed 4 % HCD and control diet, followed by 3 days fasting. **G)** LipoGlo-Counting for measuring total ApoB-LP in homogenized liver, gut, or muscle tissues of fasted fish after being fed the control diet or 4 % HCD (mean +/- SD, robust Two-Way ANOVA with Games-Howell tests). **H)** Quantification of major lipid classes in postprandial and fasted zebrafish plasma using HPLC-CAD platform. Zebrafish that were identified to carry the “fatty liver” phenotype are indicated as white dots on the bar graphs (mean +/- SD, robust Two-Way ANOVA with Games-Howell tests). * *p* < 0.05, ** *p* < 0.001, # *p* = 0.07.

A previously established HPLC method was used to detect and quantify the composition of major lipids in plasma or tissues of zebrafish (31). Total lipids of zebrafish are separated into three major groups by HPLC, which are 1) free fatty acids (FFA) and lysophospholipids (lyso-PL), 2) phospholipids and cholesterol, and 3) triacylglycerol and CE (Figure 1 B). To further identify each individual peak, we used the TLC method (33) to separate PC, PE, TG, and CE. We then ran each lipid class on HPLC separately (Supplementary Figure 1). By summarizing all identified peaks of each lipid class, we have found that fasted HCD females had significantly higher (*p* < 0.05) plasma levels of phospholipids, cholesterol, and CE as compared to fasted control females, while only CE was significantly different between males fed HCD and the control diet (Figure 4 C). In addition, females fed HCD and then fasted likely had higher (*p* = 0.07) plasma FFA and lyso-PL levels compared to females which fed the control diet before fasting (Figure 4 C).

Like in larvae, feeding 4 % HCD followed by fasting also resulted in a fatty liver phenotype in adult fish. After fasting for 3 days, 30 % of the female fed HCD developed an opaque, white liver, suggesting high levels of hepatic steatosis (Figure 4 D), while no fatty liver was observed in male fish nor in fish fed the control diet. Feeding 4 % HCD followed by 3-day fasting also increased ApoB-LP levels in female livers, however, no differences were observed in the intestine or muscle (Figure 4 E). By using HPLC, we found females fed HCD before fasting have higher TG (*p* = 0.05) and CE (*p* < 0.001) levels than females fed the control diet before fasting (Figure 4 F and Supplementary Figure 2). More interestingly, the females with fatty liver had double of amount of TG than fish with normal livers (2.8 *vs.* 1.4 pA/mg of protein), while no differences on CE was observed between fatty liver and normal liver female (Figure 4 F). In postprandial females, no difference was observed for the lipids in liver between fish fed the control and HCD diet (Figure 4 F). Feeding of HCD also had no effect on the major lipids in male livers (Figure 4 F).

### 3.5 The fasting-induced hepatic steatosis is associated with upregulated lipogenesis gene in adipocytes

To understand the molecular mechanisms underlying the HCD-induced liver steatosis, and why there was so much additional TG found when only dietary cholesterol was increased, we sequenced RNA from livers of fasted females and males fed either 4 % HCD or control diet. Although dramatic physiological and biochemical differences were observed between the livers of the fish fed HCD and the control diet, only a few differentially expressed genes (DEG = 18 in females and 12 in males, *q* < 0.05 & |log2FC| > 1) were identified (Figure 5A). None of the DEGs were involved in TG synthesis pathways or fatty acid oxidation pathways. Nine out of 12 down-regulated DEGs in HCD females were involved in cholesterol biosynthesis pathways, which was correlated with higher CE content in females fed HCD than control diets (Figure 4D). Other down-regulated DEGs included the *tyrosine aminotransferase* (*tat*) gene, which is involved in catalyzing the breakdown of L-tyrosine into p-hydroxyphenylpyruvate (44), and *cytidine monophospho-N-acetylneuraminic acid hydroxylase* (*cmah*), which hydroxylates N-acetylneuraminic acid (Neu5Ac), producing N-glycolylneuraminic acid (Neu5Gc) (45). Only 6 up-regulated DEGs were identified in females fed the HCD compared to the control diet. These DEGs include 2 genes for circadian regulation (*RAR-related orphan receptor C* (*rorcb*) and *cryptochrome circadian regulator 4* (*cry4*)). Other 3 DEGs were *prolyl 4-hydroxylase subunit alpha 1* (*p4ha1b*), which encodes a key enzyme in collagen synthesis, *phosphorylase B kinase gamma catalytic chain* (*phkg1b*) which is involved in glycogenolysis, and the *synaptotagmin 5a* (*syt5a*) gene involved in the cellular response to calcium ions. All 12 DEGs were downregulated in HCD males compared to the control males (Figure 5 A). Half of these DEGs were *vitellogenin* (*vtg*) genes which are expressed at modest levels in male zebrafish (despite their well-described role in female lipid transport to the ovary) and may function as an additional lipid and protein transporters (19). Other interesting down-regulated DEG includes *3-hydroxy-3-methylglutaryl coenzyme A reductase* (*hmgcra*) which encodes the rate-controlling enzyme in cholesterol biosynthesis (46). Differential expression analysis was also done between sexes. In general, 2201 DEGs (*q* < 0.05 & |Log2FC| > 1) were identified between HCD females and HCD males, while 1946 DEGs were found between females and males under the control diet (Supplementary Figure 7). To understand transcriptional regulation differences between males and females in general, we ran KEGG enrichment analysis on shared 1358 DEGs between the two sexes (Supplementary Figure 7). The 891 up-regulated DEGs in females were enriched in lipogenesis, including the fatty acid synthesis and steroid biosynthesis pathway (Supplementary Figure 7). On the other hand, the 467 down-regulated DEGs in females were enriched in degradation pathways, including the PPAR signaling pathway and fatty acid degradation.

**Figure 5.**
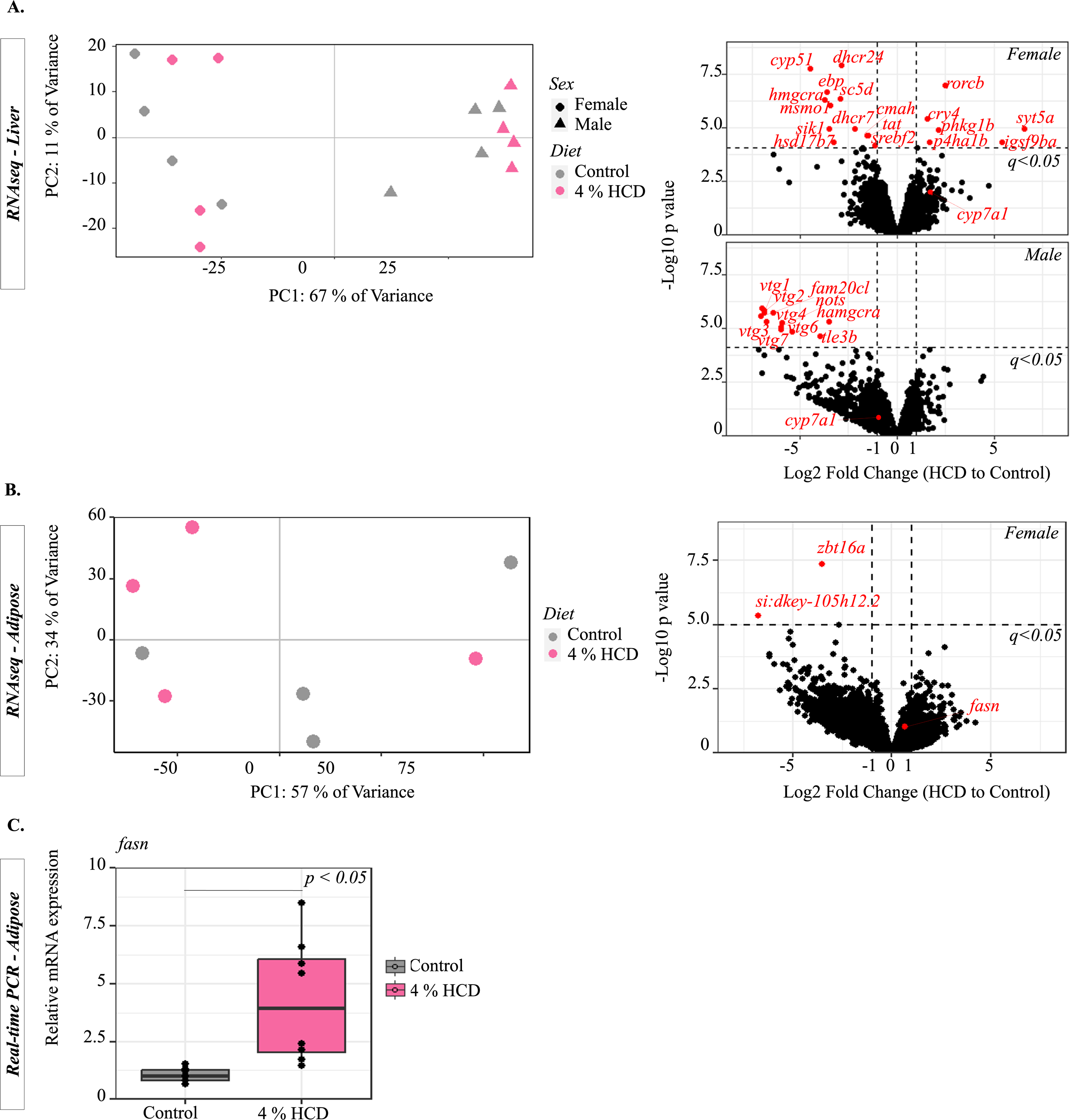
Gene expression analysis on adipose and liver of zebrafish fed 4 % HCD and control diet followed by 3 days of fasting. **A)** Transcriptomic (RNAseq) analysis on the liver of fish fed 4 % HCD compared to the control diet. The left panel shows the score plot of PCA on log2 count per million (CPM) of the top 500 most variant genes across all 16 fish. The right panel shows DEGs (*q* < 0.05 and |Log2FC| > 1) between the livers of fish of each gender fed 4 % HCD compared to the control diet. **B)** Transcriptomic analysis on the adipose tissue of fish fed 4 % HCD compared to the control diet. The left panel shows the score plot of PCA across all samples (4 individuals × 2 diets). The right panel shows DEGs between the livers of fish fed 4 % HCD and the control diet. **C)** RT-PCR measurement of *fasn* gene expression (relative to control) in adipose tissue. Each diet group contains 8 individual ish samples (n = 8) collected from 2 independent experiments (Welch t-test).

In vertebrates, fasting liberates FFA from adipocytes that are ultimately delivered to the liver, where they are essential for the synthesis of fuels needed to maintain organ function. Therefore, we measured expression of lipogenesis genes in female adipocytes that were fed either 4 % HCD or the control diet followed by a 3 day fast. The qPCR result showed that the key lipogenesis gene, *fatty acid synthase* (*fasn*) was significantly (*p* < 0.05, n = 8 fish per group) upregulated in fish fed HCD as compared to the control diet (Figure 5 C). To further investigate the gene regulatory network differences, we performed RNA-seq on abdominal visceral adipose tissue (40) of fish fed 4 % HCD and the control diet, followed by fasting for 3 days. Only two differential expressed genes (DEG, *q* < 0.05 & |log2FC| > 1) were identified, which included the *zinc finger and BTB domain-containing protein 16* (*zbt16a*) gene and an uncharacterized *si:dkey-105h12.2* gene. The expression of *fasn* gene was generally higher in fish fed 4 % HCD than control diet, though the difference was not statistically significant (*q* > 0.05, Supplementary Figure 8).

Since bile salt production can be used to reduce systemic cholesterol levels in vertebrates (5), the effect of the HCD on bile production was examined. The previously established *fabp6:GFP* reporter line expresses green fluorescent protein (GFP) in the ileal epithelium driven by a 1.7-kb zebrafish *fabp6* promoter fragment (30). This reporter line can be used to monitor bile salt signaling in zebrafish because bile salts bind and activate farnesoid X receptor (FXR) which in required to drive expression of *fabp6:GFP* (30). Indeed, we observed significantly (*p* < 0.05) higher GFP fluorescence in fish fed % HCD as compared to the control diet, which suggested that HCD promotes bile signaling (Supplementary Figure 9). This HCD-induced fluorescent difference was attenuated in animals lacking the bile synthesis gene *cyp7a1* (Supplementary Figure 9).

## 4 Discussion

Excess dietary cholesterol has been associated with various lipid disorders, including hypercholesterolemia (47), non-alcoholic fatty liver disease (NAFLD) (8, 48–50), obesity, and diabetes (51, 52). This study describes a validated and reproducible HCD protocol for zebrafish that produced phenotypes consistent with hypercholesterolemia and NAFLD. Compared to the previously established HCD method, which mixes cholesterol and a basal fish diet in diethyl ether (11, 20, 53), our improved HCD method 1) has a more user-friendly detailed protocol where its stability, reproducibility, and fish palatability were validated, 2) improved the previously described protocol (20) by using a less hazardous solvent ethanol instead of diethyl ether, and 3) has been validated with one of the most widely used commercial zebrafish diets (Gemma), of which we determined the lipid content (low TG and high phospholipids) (Figure 1 B). Zebrafish fed our HCD diet developed hypercholesterolemia, as indicated by a doubling of plasma cholesterol and ApoB-LP levels (Figure 4). In humans, hypercholesterolemia is closely associated with atherosclerosis and high circulating LDL. Although we did not observe atherosclerotic plaques as others have previously described (20, 54), the elevated plasma cholesterol levels and atherogenic ApoB-LP justify the use of the zebrafish as a model for identifying factors that influence the development of human cardiovascular disease.

Another striking observation was that under the HCD, the zebrafish developed severe hepatic steatosis, as indicated by liver opacity (Figure 3). Unlike previous studies which identified hepatic steatosis only using Oil Red O or Nile Red staining (26, 55), our HCD has introduced strong hepatic lipid droplet accumulation which scatters transmitted light (56) and produces the readily visible opaque liver phenotype under brightfield microscopy. Given that the zebrafish larvae are well-suited for high-throughput drug screening, this opaque liver phenotype provides a tractable opportunity to identify small molecules that attenuate liver steatosis.

In our studies, the development of liver steatosis was strongly associated with fasting, suggesting that high dietary cholesterol and fasting interact synergistically to induce the metabolic and hepatic features that lead to steatosis (Figure 3 C and D). In mammals, fasting is characterized by the breakdown of TG stored in white adipose tissue into FFA, which then circulate in the plasma and are utilized by the liver and other organs to maintain energy homeostasis (41). If the FFA supply is overloaded or if the liver β-oxidation pathway is inhibited, liver FFA will be re-esterified into TG and accumulate in cytoplasmic lipid droplets (57). Transcriptomic analysis of animals subject to the HCD and fasting found no evidence of attenuated β-oxidation gene expression. However, HCD may elevate FFA levels in plasma, though this effect did not reach significance (*p* = 0.07). These findings suggest that HCD increased liver FFA uptake during fasting and likely produced the observed hepatic steatosis.

One paradox of our study was that hepatic steatosis (increased liver TG) resulted from feeding only excess dietary cholesterol (Figure 4). The necessity of fasting in addition to the high dietary cholesterol, further suggested that the interaction between adipocytes and liver contributed to the development of hepatic steatosis. Consistent with this hypothesis, HCD increased adipocyte expression of fatty acid synthase (*fasn),* a key lipogenic gene. In prior studies, knockdown of Promyelocytic leukemia zinc finger protein (Plzf, encoded by *zbtb16* gene) was associated with increased adipogenesis and lipid accumulation in mouse adipose tissues (58). Our transcriptomic data showed that the *zbtb16a* gene was the most down-regulated gene (Log2 fold change = -3.5, *q* < 0.001) in white adipose tissue in HCD-fed fish. Increased lipogenesis might provide HCD fish with a larger TG pool in adipocytes which supplies liver FFA during fasting. Mouse and primate studies in which increased dietary cholesterol was associated with adipocyte hypertrophy and increased free cholesterol FC content in visceral adipose tissue (51, 59), which are consistent with our zebrafish data from animals fed an HCD. Adipocytes are also known to be major vertebrate FC depot (48). Studies have also shown that the activation of lipolysis during fasting decreases the cellular level of TG in proportional to the cellular FC levels (59, 60), which supports our observation that fish fed HCD had increased plasma FC as well as FFA levels during fasting (Figure 4 C). We also observed increased FC in proportion to TG levels in fasted HCD fish (Figure 4 E). As intracellular FC is primarily stored in the lipid droplet surface layer (61), it is likely that hepatocyte lipid droplets need to maintain a FC:TG ratio and synthesize extra TG to balance the excess imported FC during fasting.

By comparing hepatic transcriptomes between HCD and control fish, we did not identify DEGs in *de-novo* lipogenesis, β-oxidation, or VLDL synthesis pathways, suggesting that these pathways had little contribution to the hepatic steatosis. This was contradictory to previous studies, which identified liver-specific down-regulation of β-oxidation gene *Cpt-1a*, VLDL synthesis gene *Mttp*, and lipogenesis gene *Fasn* of mice fed HCD (62, 63). One possible explanation for this is that in these studies, mice were exposed to an HCD for a much longer period (30 weeks) than zebrafish (9 days) which may alter liver lipid metabolism more severely. Prolonged fasting also causes liver steatosis in animals under a normal diet, which is suggested to be caused by reduced liver VLDL production during fasting that limits TG efflux (64). However, the VLDL production is less likely the limiting factor for the zebrafish HCD-induced liver steatosis we observed since we found that plasma VLDL levels doubled in fish fed the HCD (Figure 4 A). The fasting-induced fatty liver phenotype was only observed in female zebrafish, which makes sex another critical factor for the development of liver steatosis. This correlates with human studies in which women had higher lipolysis and plasma FFA than men after fasting (65). RNA-seq results indicated higher expression of lipid synthetic and transport genes, along with lower expression of fatty acid β-oxidation genes in females, which again suggests that the sex difference in lipid metabolism contributed to the sexual dimorphism in fasting induced liver steatosis. The early stages of non-alcoholic fatty liver disease (NAFLD) are characterized by simple hepatic steatosis, which can progress to a more aggressive stage called non-alcoholic steatohepatitis (NASH) that can then develop inflammation with or without fibrosis (66). As fibrosis develops, NASH can progress to cirrhosis, the end-stage of NAFLD. Although the molecular mechanism underlying the progress of NAFLD remains unclear, many studies have suggested that hepatic FC is critical for the development and progression to NAFLD (6, 8, 59). Our study has identified clear hepatic steatosis and higher levels of FC and TG in zebrafish under HCD. This suggests that the fish develop similar liver lipid phenotypes as those found in early-stage human NAFLD (Figure 4). Additionally, we have identified the upregulation of the *p4ha1* gene, which encodes a key enzyme in collagen synthesis (67). As liver fibrosis is characterized by excess deposition of collagen, the upregulation of *p4ha1* suggests that liver fibrosis may be induced in zebrafish under HCD. The zebrafish HCD model may have the potential to develop NASH by prolonging the fasting period or introducing additional dietary TG. Further, the inhibition of P4ha1 might decrease collagen production and work as a therapeutic for treating liver fibrosis (67–69).

In conclusion, this well validated HCD zebrafish model increases ApoB-LP levels and results in liver steatosis, which makes it tractable for studying the underlying biology of hypercholesterolemia and NAFLD. In addition, zebrafish fed extra dietary cholesterol developed hepatic steatosis as indicated by increasing TG levels, likely caused by enhanced adipose lipolysis and increased plasma FFA that combine to overload the liver during fasting.

## Data Availability

Raw sequences are publicly available on Sequence Read Archive (SRA) under accession number PRJNA998935.

## Acknowledgements

We thank Dr. Meredith H. Wilson for reviewing the final manuscript. We gratefully acknowledge Julia Baer and Mackenzie Klemek for fish husbandry.

