## Supplementary Table and Figures for "A high-cholesterol zebrafish diet promotes hypercholesterolemia and fasting-associated liver triglycerides accumulation"

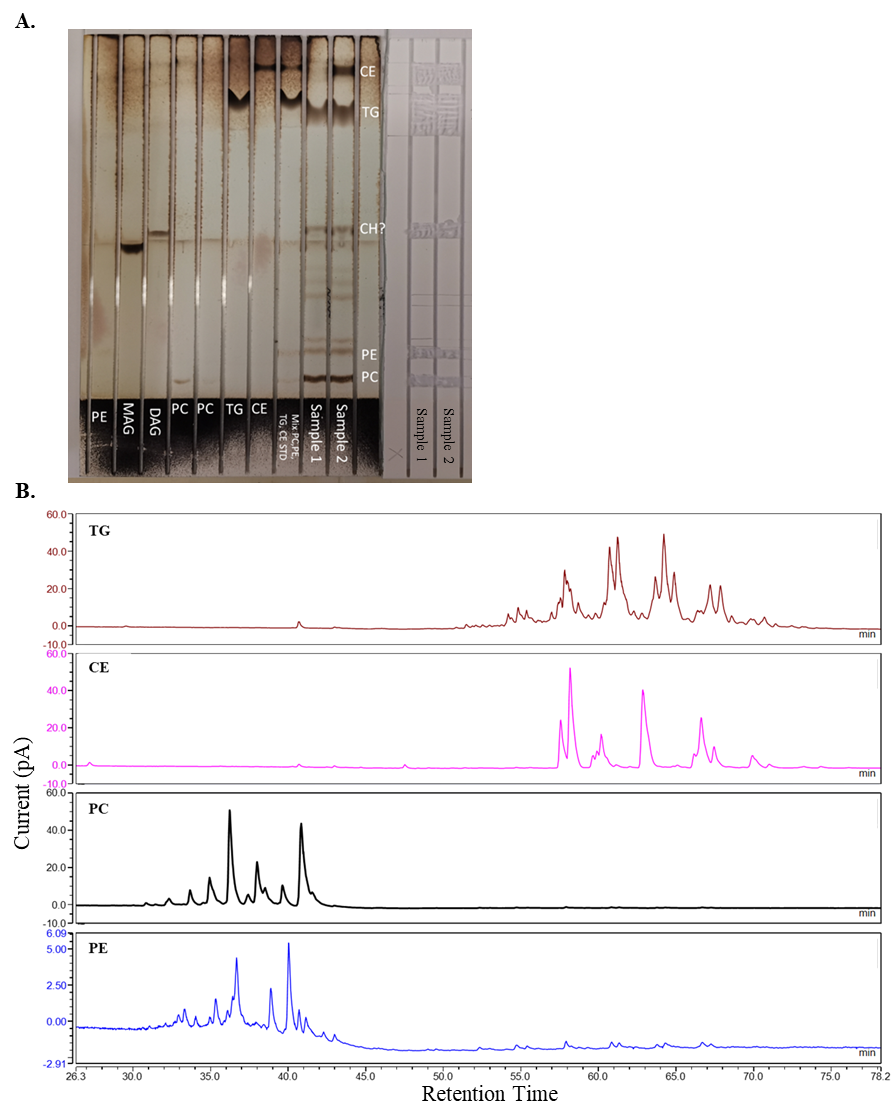

**Supplementary Figure 1.** **A)** A TLC plate which shows charred lipid standards and samples on the left and uncharred samples for HPLC analysis. **B)** Chromatogram figures for lipid classes separated from total lipids of zebrafish adult liver. Thin-layer chromatography (TLC) technology separated different lipid classes before running as separate samples on HPLC.

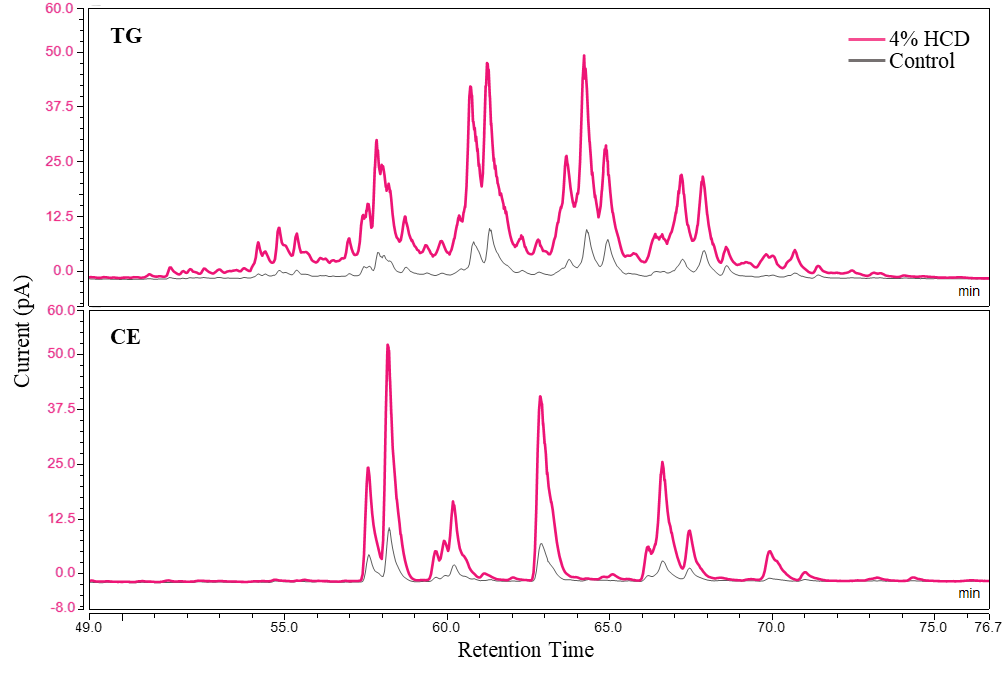

**Supplementary Figure 2. Chromatogram figures for separated TG and CE from total lipids of zebrafish liver.** Fish were fed either 4 % HCD or the control diet for 2 weeks, followed by a 3-day fast. TLC technology separated different lipid classes before running them as separate samples on HPLC.

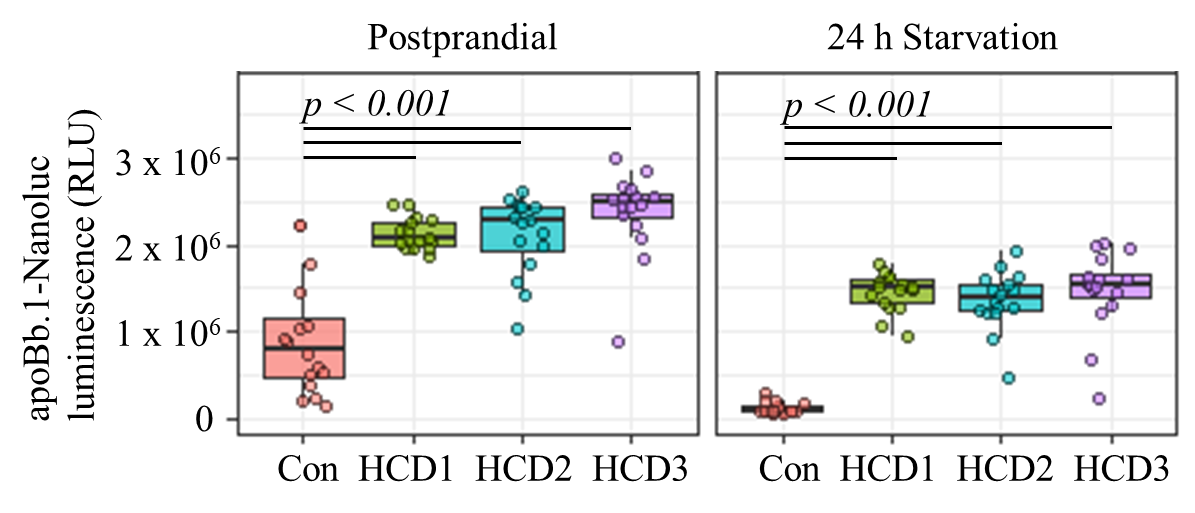

**Supplementary Figure 3. Effect of HCD made by 3 different methods on total ApoB-LP levels (LipoGlo-counting) of 14 dpf fish.** Control diet: standard GEMMA 75 diet treated with 100 % ethanol. HCD1: our standard HCD protocol (Figure 1 A), except that the mixture was vacuum treated before drying. HCD2: our standard HCD protocol (Figure 1A). HCD3: previously published HCD protocol which used ether to deliver cholesterol to GEMMA diet (20). Fifteen fish were used in each dietary group (n = 15). Two-way robust ANOVA and Games–Howell test were used for all samples.

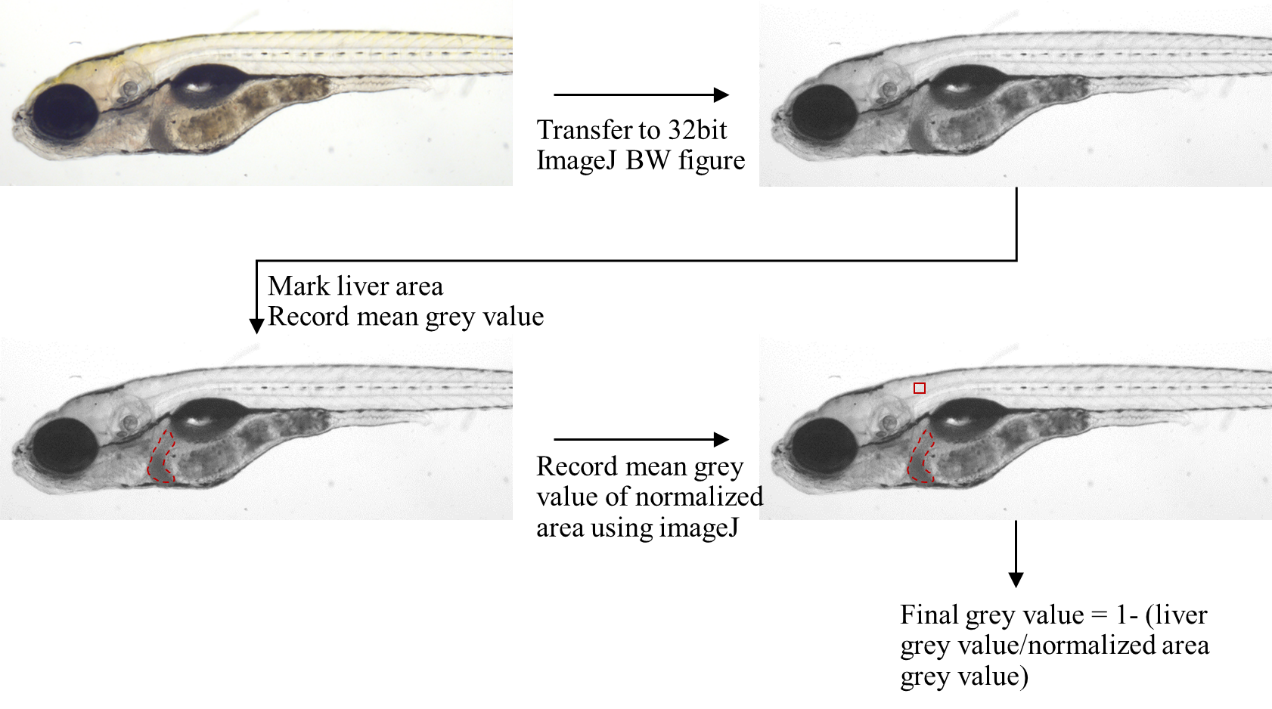

**Supplementary Figure 4. Method for measuring opacity of the liver.** Images were first transferred to 32-bit ImageJ black-and-white figures. Liver areas were marked, and mean grey values were measured using ImageJ. A 50x50 square was drawn in the fixed area in the muscle tissue near the liver for normalization. All images from the same day were taken under the same light intensity.

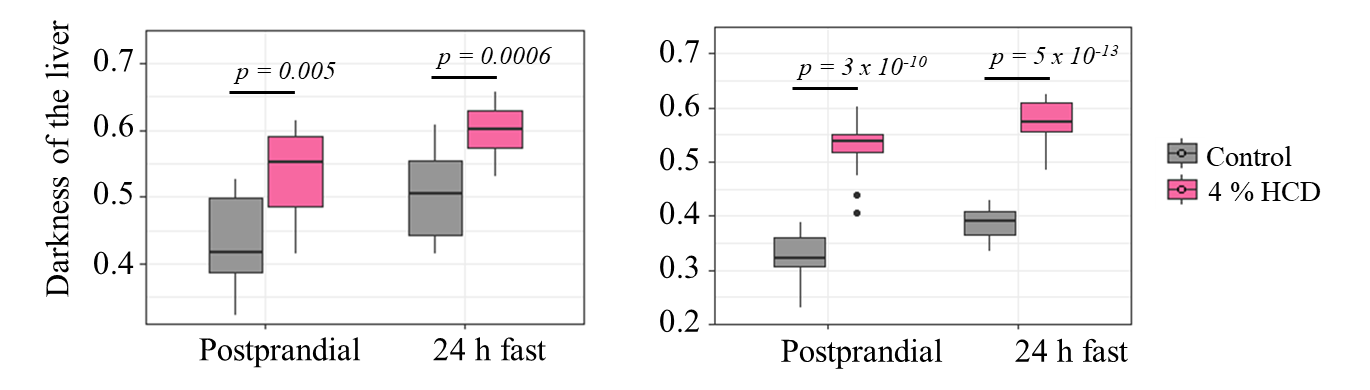

**Supplementary Figure 5**. **Effect of fasting on liver opacity between fish stocks.** Fish were fed either the control diet or 4 % HCD from 6 dpf to 13 dpf, followed by either continued feeding or fasting for 24h. Two-way robust ANOVA and Games–Howell test were used.

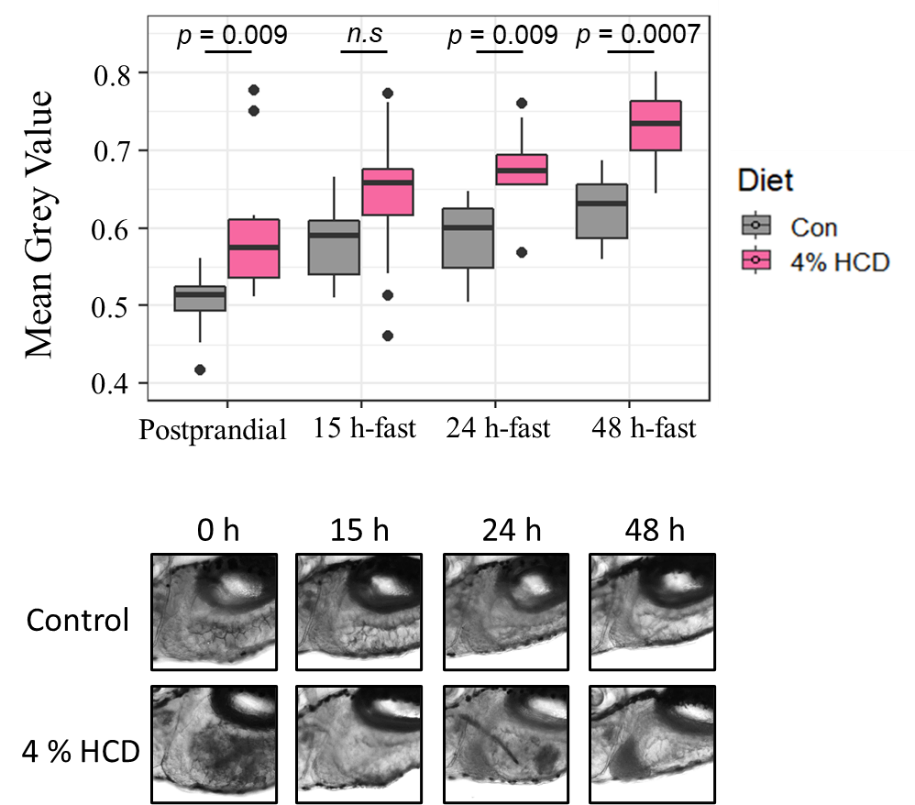

**Supplementary Figure 6.** **Effect of fasting period on liver opacity.** Fish were fed either the control diet or 4 % HCD from 6 dpf to 13 dpf, followed by fasting. Individual fish were tracked and imaged at 0 h, 15 h, 24 h, and 48 h after fasting (n = 6 per each diet group). The top panel is merged data from 2 independent experiments. Two-way robust ANOVA and Games–Howell test were used for all samples. The bottom panel is representative of individual images following fasting.

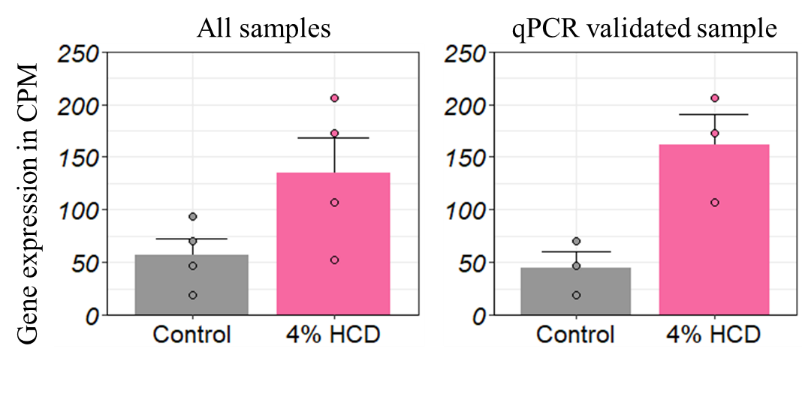

**Supplementary Figure 7. Expression of *fasn* gene between fish fed control diet and 4% HCD.** The left figure showed the expression of all 4 samples in each group used for RNA-seq analysis. The expression of gene was showed in count per million (CPM) value which is raw counts normalized by total counts of each sample. The right figure showed the expression of 3 samples in each group which were used for qPCR. Statistical test was done by using R package edgeR, and the expression of *fasn* gene showed no difference (*q* < 0.05) between fish fed 4% HCD and control diet.

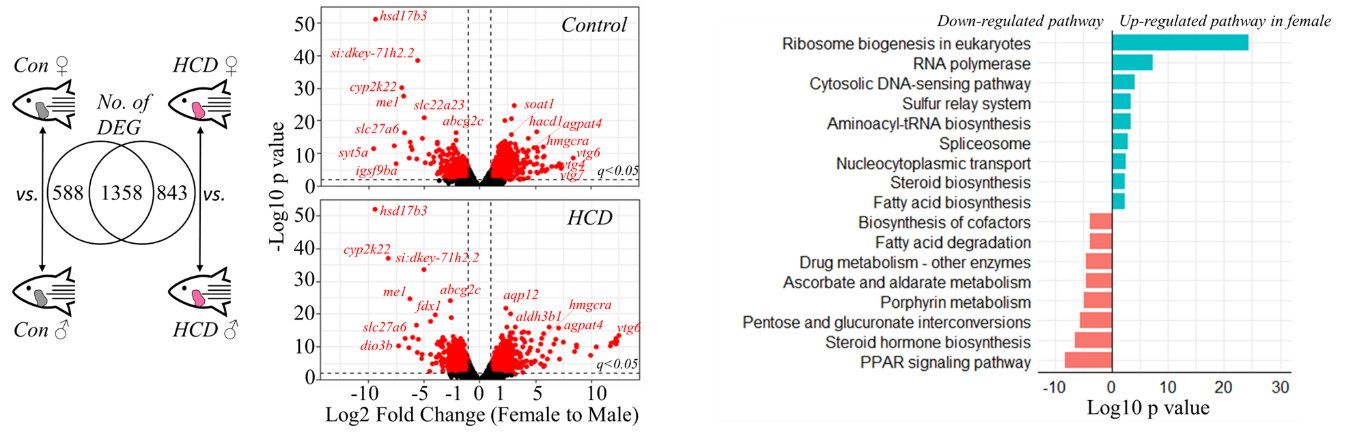

**Supplementary Figure 8.** **DEGs between the liver of female fish compared to male fish.** KEGG enrichment shows the top significant enriched pathway for the common 1358 DEGs between male and female fish regardless of dietary treatment.

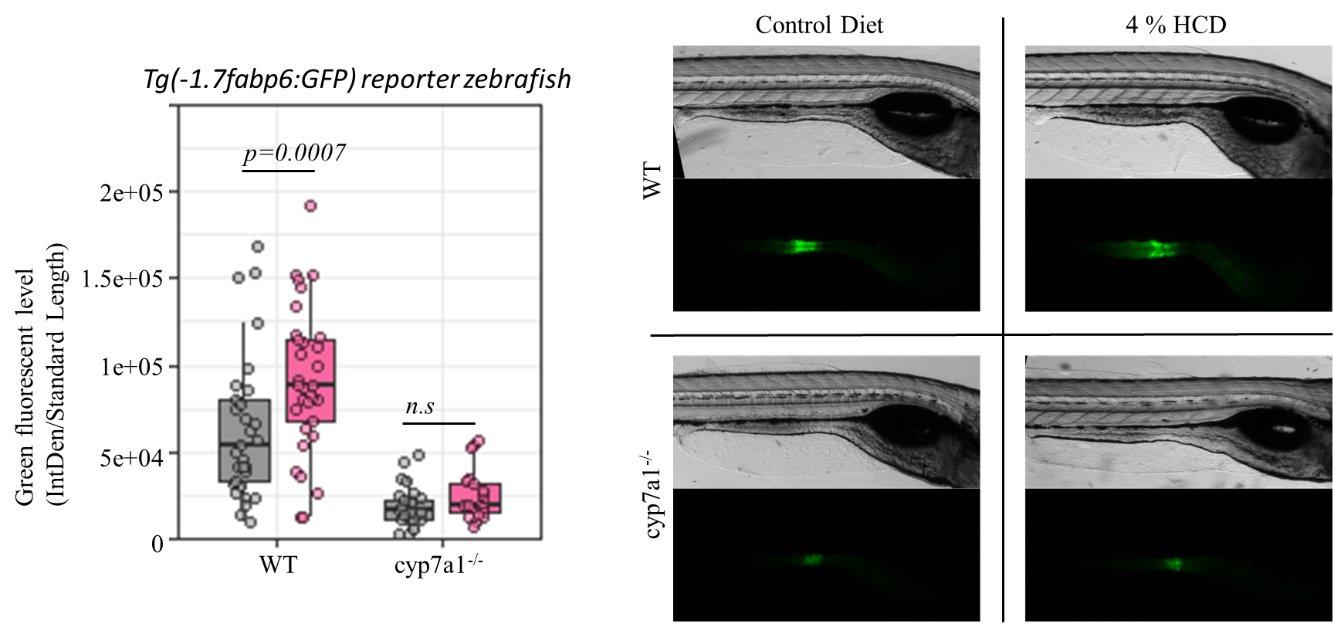
**Supplementary Figure 9. Effect of 4 % HCD on bile acid signaling in zebrafish.** The left panel shows total green fluorescent protein fluorescence level in *Tg(-1.7fabp6:GFP)* reporter zebrafish with or without *cyp7a1* mutation fed either 4 % or the control diet. As *fabp6* is a Fxr target, regulated by bile salts in zebrafish, this result suggests that our HCD could be used for modulating bile salt signaling in zebrafish. Two-way robust ANOVA and Games–Howell test were used for all samples.
